# Pathway metabolite ratios reveal distinctive glutamine metabolism in a subset of proliferating cells

**DOI:** 10.1101/2024.02.18.580900

**Authors:** Nancy T Santiappillai, Yue Cao, Mariam F Hakeem-Sanni, Jean Yang, Lake-Ee Quek, Andrew J Hoy

**Affiliations:** The University of Sydney, Charles Perkins Centre, School of Medical Sciences, Sydney, New South Wales, 2006, Australia; The University of Sydney, Charles Perkins Centre, School of Mathematics and Statistics, Sydney, New South Wales, 2006, Australia; The University of Sydney, Sydney Precision Data Science Centre, Sydney, New South Wales, 2006, Australia

**Keywords:** metabolomics, metabolic pathways, cell lines, cancer, glutamine metabolism, glucose metabolism

## Abstract

Large-scale metabolomic analyses of pan-cancer cell line panels have provided significant insights into the relationships between metabolism and cancer cell biology. Here, we took a pathway-centric approach by transforming targeted metabolomic data into ratios to study associations between reactant and product metabolites in a panel of cancer and non-cancer cell lines. We identified five clusters of cells from various tissue origins. Of these, cells in Cluster 4 had high ratios of TCA cycle metabolites relative to pyruvate, produced more lactate yet consumed less glucose and glutamine, and greater OXPHOS activity compared to Cluster 3 cells with low TCA cycle metabolite ratios. This was due to more glutamine cataplerotic efflux and not glycolysis in cells of Cluster 4. *In silico* analyses of loss-of-function and drug sensitivity screens showed that Cluster 4 cells were more susceptible to gene deletion and drug targeting of lactate and glutamine metabolism, and OXPHOS than cells in Cluster 3. Our results highlight the potential of pathway-centric approaches to reveal new aspects of cellular metabolism from metabolomic data.

## INTRODUCTION

Cancer cells are characterized by dynamic plasticity of nutrient utilization that supports tumor growth and survival (Altea-Manzano *et al*, 2020; Fendt *et al*, 2020; Pavlova *et al*, 2022). Our understanding of the role of metabolism in cancer cell biology has predominantly arisen from studies taking cancer type-specific and/or metabolic pathway-focused approaches (for example, Hensley *et al*, 2016, Kamphorst *et al*, 2015, and Wang *et al*, 2023).

More recently, several publications have reported the outcomes of high-throughput metabolomics using pan-cancer cell line panels, such as the NCI-60 (60 cell lines, Jain *et al*, 2012; Ortmayr *et al*, 2019) and CCLE panels (928 cell lines, Li *et al*, 2019), CAMP (988 tissue samples, Benedetti *et al*, 2023), and other panels containing 180 cell lines (Cherkaoui *et al*, 2022), and 173 cell lines (Shorthouse *et al*, 2022). These studies primarily aimed to identify links between cancer cell metabolic phenotypes and transcriptional regulation (Benedetti *et al*., 2023; Ortmayr *et al*., 2019), or genetic alterations and dependencies (Li *et al*., 2019; Mullen & Singh, 2023) that were associated with drug-sensitivities (Shorthouse *et al*., 2022). Notably, Cherkaoui and colleagues (2022) took a top-down approach by clustering the metabolome acquired by untargeted metabolomics across 49 KEGG metabolic pathways of 180 cancer cells. Pathway activity was determined for each cell line, and they identified only two clusters that were defined by either high carbohydrate metabolic activity or high aerobic mitochondrial activity, that was associated with epithelial or mesenchymal status, respectively (Cherkaoui *et al*., 2022).

Here, we took a different approach to identify common signatures based upon high-flux metabolic pathways only in a smaller pan-cancer panel of proliferating cells. Based on metabolite levels, we initially identified four clusters of cells, but this approach failed to provide insights into pathway differences. To overcome this issue, we introduce a conceptual innovation to transform our metabolite data into pathway-centric ratios that resulted in the formation of five clusters of cells that displayed different ratios of metabolites of glycolysis, pentose phosphate pathway, pyruvate-TCA cycle, proline metabolism, serine metabolism, glutamine metabolism, and methionine metabolism. Of these five clusters, we used a combination of techniques to show that cells in Cluster 4 had higher ratios of TCA cycle metabolites when normalized to pyruvate and produced more lactate, despite lower glucose and glutamine consumption, and greater OXPHOS activity than Cluster 3 with low TCA cycle metabolite ratios. These differences were, in part, explained by increased glutamine cataplerotic efflux and glutaminolysis. These phenotypes were supported by *in silico* analyses of pan-cancer loss-of-function and drug sensitivity screens to show that cells in Cluster 4 were more susceptible to gene deletion and drug targeting of lactate and glutamine metabolism, and OXPHOS compared to the cells in Cluster 3. These results highlight the benefit of converting metabolite levels into pathway-based ratios as a starting point for gaining insights into cellular metabolic activity.

## RESULTS

### Targeted metabolomic profiling of high flux pathways in cell lines from 11 tissue origins

We quantified the metabolite levels of high flux pathways, including central carbon (glycolysis, TCA cycle, pentose phosphate pathway) and amino acid metabolic pathways, in 57 adherent cell lines (49 tumor- and 8 normal epithelial-derived) from 11 cancer types cultured in basal media conditions (Appendix Table 1). Samples were generated in triplicate across 6 batches, including control cell lines for batch correction, ensuring consistency throughout the complete data set (Fig 1A). K-means algorithms with Pearson’s correlation of the batch-corrected dataset identified 4 distinct clusters of cells (Fig 1B), which were not a consequence of the culturing conditions, tissue type (normal epithelial vs. tumor), cancer type, tissue origin, or the mutation status of common oncogenic drivers that influence cell metabolism (Cairns *et al*, 2011; Jia *et al*, 2008; Jones & Thompson, 2009; Oermann *et al*, 2012; Vousden & Ryan, 2009) (Fig 1C).

**Figure 1:**
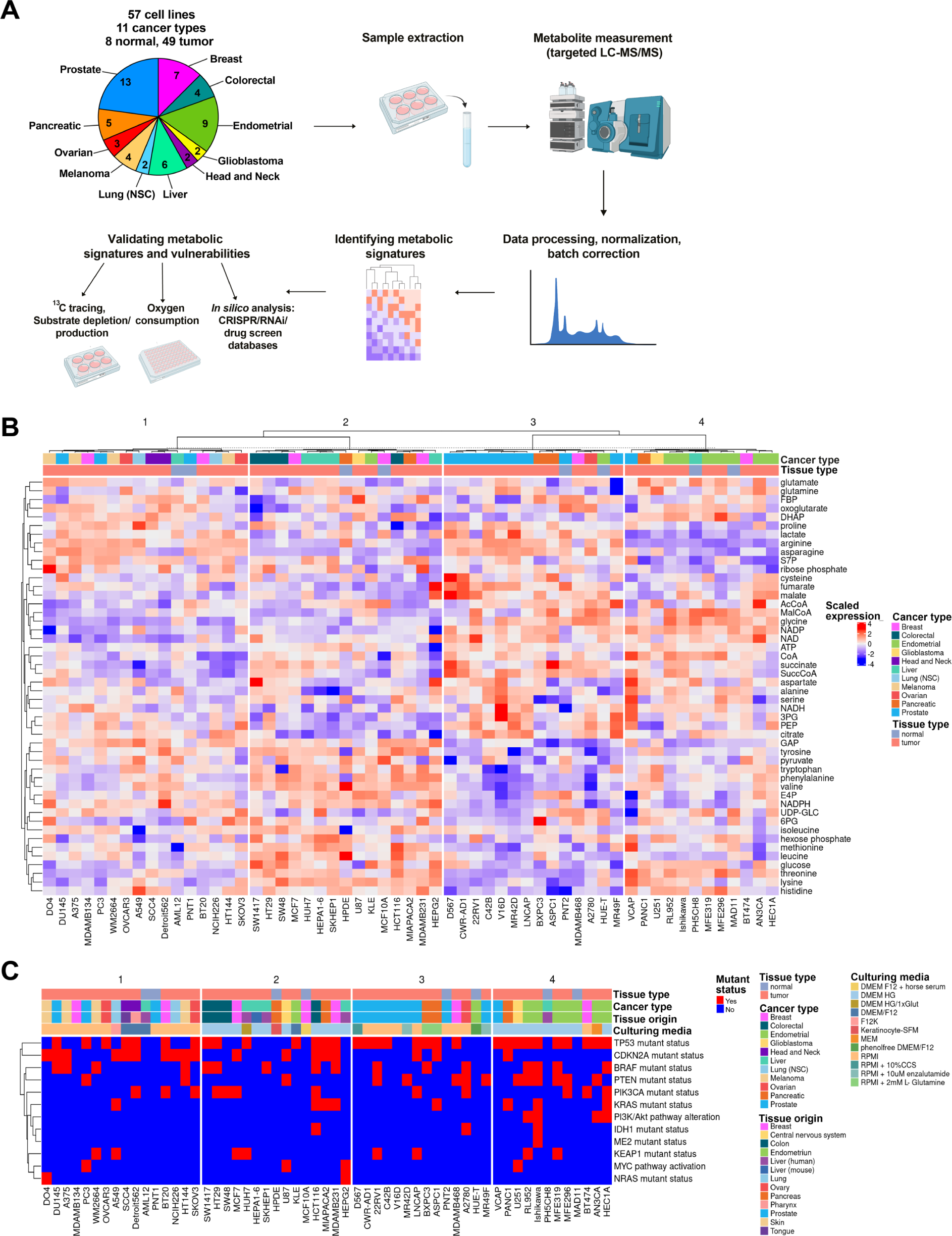
The metabolome landscape of cells from 11 tissue origin sites. A. Schematic of the workflow for targeted metabolomics profiling of 57 cell lines to identify and validate metabolic signatures. B. Heatmap of scaled metabolite expression across central carbon and amino acid metabolism within the cell line panel (n=57). Cell lines color coded by cancer type and tissue type. C. Clusters of cell lines from (B) appended with color coded legends for mutant status of common oncogenic drivers, culturing media conditions, tissue origins, cancer type, and tissue types.

All clusters contained tumor and normal cell lines from different tissue origins (Fig 1B). However, there were some instances where cell lines from the same tissue origin clustered together, such as Cluster 3 that was enriched with prostate cancer cells and Cluster 4 with endometrial cancer cells (Fig 1B), which has been observed in another pan-cancer metabolome study (Shorthouse *et al*., 2022). Despite the identification of heterogeneous clusters of cells based upon the levels of metabolites of high flux pathways, there was no clear organization of these metabolites into pathways (Fig 1B), likely because strong metabolite interactions are often localized at the reaction level (Benedetti *et al*., 2023), that could underpin a testable hypothesis centered on differences and similarities of pathway activity.

### Analyses of pathway-centric metabolite ratios uncover distinctive metabolic signatures

Next, we took a physiological-based approach and evaluated the hypothesis that there were differences in high-flux metabolic pathways in our panel of cancer and epithelial cells. To achieve this, we transformed our metabolomic data by calculating the ratios between an upstream precursor or reactant metabolite (applied as the denominator) and downstream pathway product metabolites (numerator) for each central carbon and major amino acid metabolism pathway, similar to Benedetti *et al*. (2023). This pathway-centric transformation was based on the idea that metabolite conversion forms a cascade, and therefore, intrinsic correlations likely exist between reactant and product that provide biologically meaningful insights into pathway activity. For example, glucose is the precursor of glycolysis, and as such, ratios of the abundance of glycolytic metabolites relative to glucose were calculated (Fig 2A).

**Figure 2:**
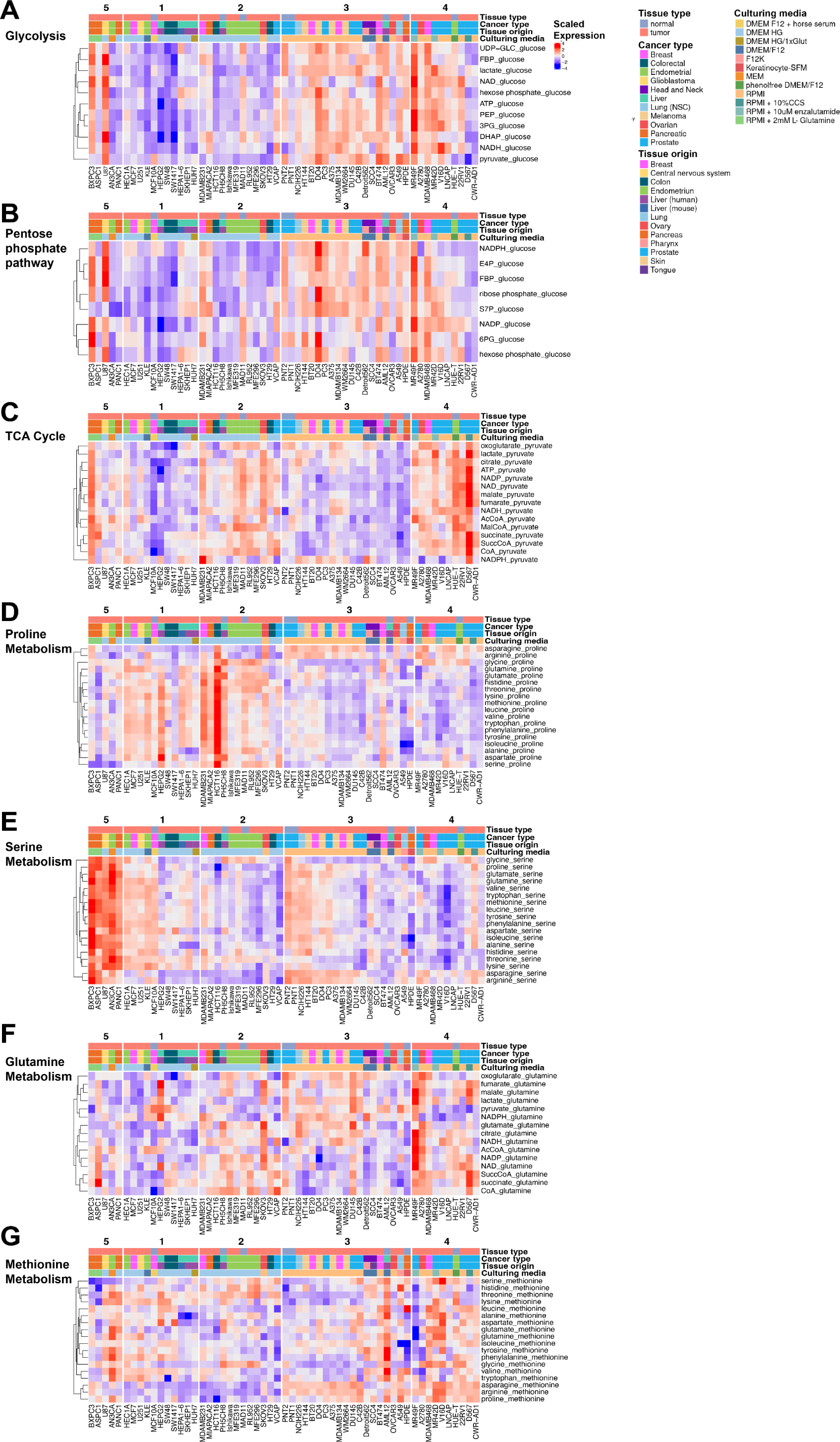
Pathway-centric metabolite ratios identifies clusters of cells. A-G. Metabolite ratio heatmaps formed by normalization of (A) glycolysis pathway and (B) pentose phosphate pathway metabolites to intracellular glucose, (C) TCA cycle pathway metabolites to pyruvate, (D) proline metabolism metabolites to proline, (E) serine metabolism metabolites to serine, (F) glutamine metabolism metabolites to glutamine, and (G) methionine metabolism to methionine. Heatmaps are slices from Appendix Figure S1.

K-means algorithms with Pearson’s correlation of the entire metabolite ratio dataset spanning seven metabolic pathways of interest (glycolysis, pentose phosphate pathway, pyruvate-TCA cycle, proline metabolism, serine metabolism, glutamine metabolism, and methionine metabolism) identified 5 distinct clusters of cells (Appendix Fig S1A). These clusters differed from what was identified using metabolite abundances alone (Fig 1) and were composed of cells from different tissue origins, tissue types, cancer types, mutation status of common oncogenic drivers, and culturing conditions (Appendix Fig S1B).

To assist in interpreting the patterns in the data, the primary heatmap (Appendix Fig S1A) was separated into individual panels with the cell clusters conserved (Fig 2). The subset of cells in Cluster 3 had greater glycolysis (Fig 2A) and pentose phosphate pathway metabolite ratios (PPP; Fig 2B) but lower TCA cycle (relative to pyruvate; Fig 2C) and proline metabolism ratios (Fig 2D) compared to Cluster 2. We identified differences in serine metabolism between cells in Clusters 4 and 5 (Fig 2E) and less striking differences in glutamine (Fig 2F) and methionine (Fig 2G) metabolism between clusters. Together, our approach of transforming metabolite levels into pathway-specific ratios identified groups of cancer cells from different tissue lineages defined by differences in high flux metabolic pathways, not evident by metabolite levels alone.

### Differences in TCA cycle metabolite to pyruvate ratios are due to glutamine and not glucose metabolism

We next addressed a significant limitation of metabolomic data. To date, irrespective of how the metabolomics data is analyzed, we often cannot infer function/flux. Here, we use our pathway-centric analyses to validate that the clusters formed (from where??) from our pathway-centric analyses exhibited functional differences. Since the TCA cycle is an essential hub where various pathways converge and is critical for energy and biomass production and cell viability (Spinelli & Haigis, 2018) (Fig 3A), we chose to contrast Cluster 3 and Cluster 4 from the identified five clusters formed in Figure 2, as they had distinctive TCA cycle metabolite levels relative to pyruvate (Fig 3B). Firstly, Cluster 4 cells had greater TCA cycle metabolite levels relative to pyruvate (Multiple unpaired t-tests, P-value<0.001), except for oxoglutarate and NADPH (Fig 3C). Differences in succinate and malate levels relative to pyruvate were also evident at the cell line level (Fig 3D).

**Figure 3:**
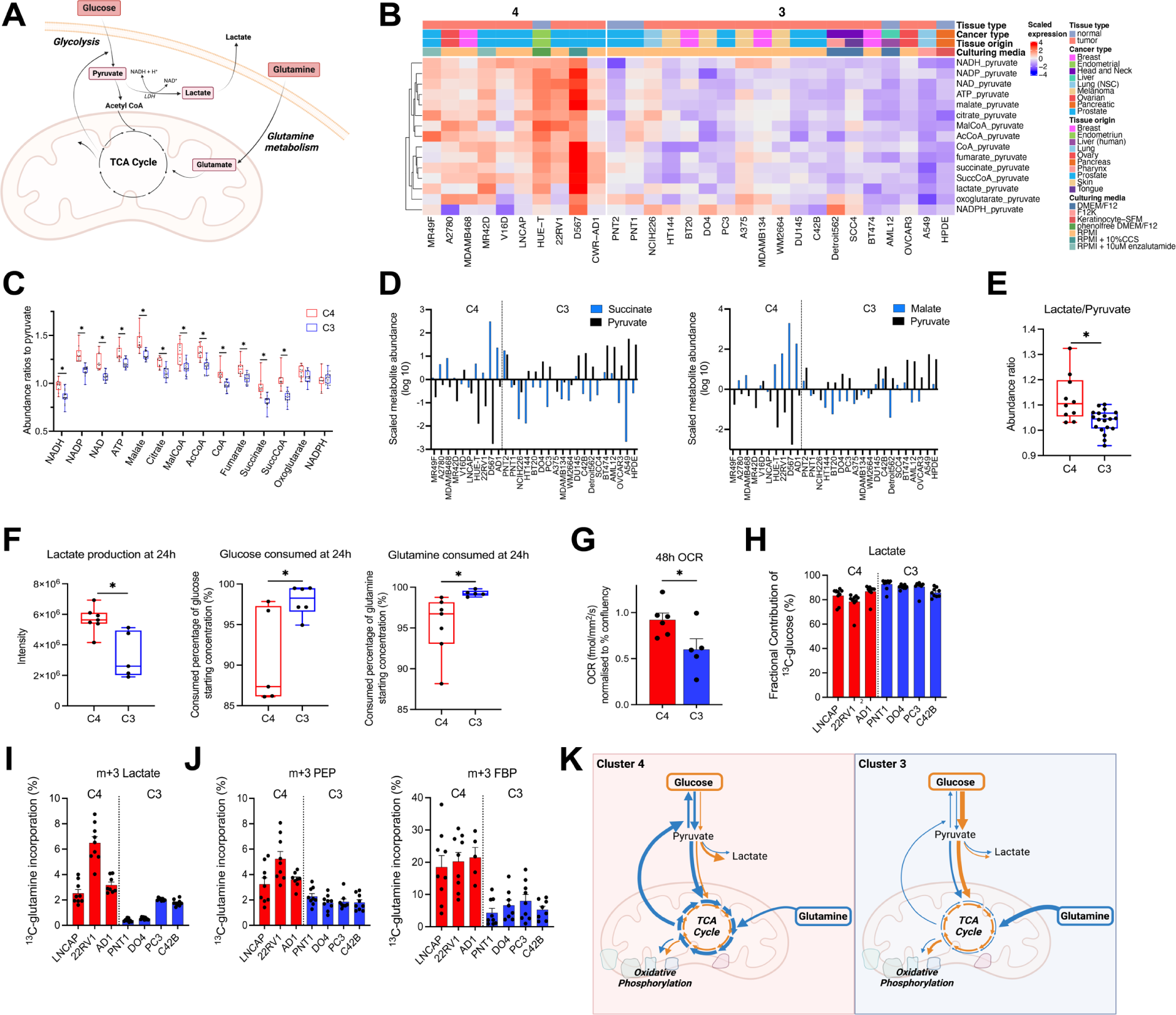
Metabolite ratio-based signatures identifies differences in glutamine metabolism between clusters. A. Schematic of glucose and glutamine sources to the TCA cycle. B. Scaled metabolite ratio heatmap of Clusters 4 and 3, derived from TCA cycle pathway metabolites calculated using the precursor pyruvate as denominator. C. Abundance ratios of TCA cycle metabolites to pyruvate (C4 n=10, C3 n=19, multiple unpaired t-tests, *p<0.05, error min to max values). D. Waterfall plots comparing the scaled metabolite abundance (log 10) of succinate and pyruvate (left), and malate and pyruvate (right) for cells of Clusters 3 and 4. E. Abundance ratio of lactate to pyruvate (error min to max values) (C4 n=10, C3 n=19). F. Lactate production intensity and glucose and glutamine consumption (consumed % of starting substrate concentrations) after 24 hours (C4 n=8, C3 n-6, error min to max values). G. Oxygen consumption rates (OCR) at 48 hours, normalized to percent confluency (C4 n=6, C3 n-5, 3 biological replicates and 4 technical replicates per cell line, mean ± standard error of the mean). H. Fractional contribution of [U-^13^C]-glucose to intracellular lactate (C4 n=3, C3 n=4, 3 biological replicates and 3 technical replicates per cell line, mean ± standard error of the mean). I. Fractional contribution of [U-^13^C]-glutamine to intracellular lactate (C4 n=3, C3 n=4, 3 biological replicates and 3 technical replicates per cell line, mean ± standard error of the mean). J. Fractional contribution of [U-^13^C]-glutamine to m+3 fructose 1,6-bisphosphate (FBP), and m+3 phosphoenolpyruvate (PEP) (C4 n=3, C3 n=4, 3 biological replicates and 3 technical replicates per cell line, mean ± standard error of the mean). K. Schematic of the C4 phenotype (increased glutamine to the TCA cycle, gluconeogenesis, and aerobic glycolysis) compared to C3 (increased glucose and glutamine consumption and glucose oxidation). Data information: (C-G) *P<0.05 vs Cluster 4 by Unpaired student’s t-test.

As pyruvate was used as the denominator to calculate TCA cycle metabolite ratios, we extended our analysis to lactate since pyruvate is converted to lactate by lactate dehydrogenase. Cells in Cluster 4 had a greater lactate-to-pyruvate ratio, a measure of the equilibrium constant of lactate dehydrogenase, than cells in Cluster 3 (Student t-test, P=0.001; Fig 3E). There was no difference in the NADH/NAD+ ratio (Appendix Fig S2A), which are co-factors of lactate dehydrogenase and can influence its activity (Luengo *et al*, 2021). Thus, the greater lactate to pyruvate ratio in Cluster 4 cells, compared to Cluster 3 cells, was unlikely driven by excess NADH relative to NAD+, but possibly due to carbon surplus in the TCA cycle.

Glycolysis and glutaminolysis both feed the TCA cycle and can produce lactate as a by-product (Smith *et al*, 2016). Next, we sought to determine whether the increased lactate production relative to pyruvate in the cells of Cluster 4 compared to Cluster 3 cells was due to greater consumption of glucose and/or glutamine. As expected, cells in Cluster 4 produced more lactate compared to those in Cluster 3 (Student t-test, P=0.006; Fig 3F), with some cell line-specific differences observed (Appendix Fig S2B). Likewise, there were cell line-specific differences in glucose and glutamine consumption over 24 hours (Appendix Fig S2B), but somewhat surprisingly, the net consumption of glucose (Student t-test, P-value=0.02) and glutamine (Student t-test, P=0.04) were lower in Cluster 4 cells (Fig 3F). In isolation, the higher lactate-to-glucose yield seen in Cluster 4 could be interpreted as higher aerobic glycolysis, whereas in Cluster 3 cells, proportionally more glucose was oxidized instead of being converted to lactate.

We postulated that Cluster 4 cells are more oxidatively competent and thus have surplus carbon that spills into lactate. To test this, we quantified the oxygen consumption rate as a readout of oxidative phosphorylation (OXPHOS) activity and, thus, TCA cycle fluxes. In line with our prediction, cells in Cluster 4 possessed greater OXPHOS activity than Cluster 3 cells (Student t-test, P=0.04; Fig 3G, Appendix Fig S3C). The more efficient respiration may correlate with our observation of more abundant TCA cycle metabolites relative to pyruvate (Fig 3C) and suggests that the increase in lactate production (Fig 3F) is a consequence of cells using pyruvate as a redox sink to regenerate NAD^+^.

To further resolve the intersection of glucose and glutamine metabolism at pyruvate, which underpins the formation of Clusters 3 and 4, we performed [U-^13^C]-glucose and [U-^13^C]-glutamine tracing experiments in a subset of cells selected from Clusters 3 and 4. As expected, lactate was produced mainly from glucose (78-92% enrichment). However, there were no differences in lactate and pyruvate enrichment between Clusters 3 and 4 (Fig 3H, Appendix Fig S2D), nor were there differences in the enrichment of TCA cycle metabolites from glucose (Appendix Fig S2E). These data, therefore, eliminate the role of glucose metabolism in explaining the higher TCA metabolite to pyruvate ratios in Cluster 4 cells compared to Cluster 3.

As such, we turned our attention to quantifying glutamine metabolism, as it is a major TCA cycle carbon source (Spinelli & Haigis, 2018), using [U-^13^C]-glutamine tracing. Our first observation was that more glutamine carbons were incorporated in lactate, via either phosphoenolpyruvate carboxykinase or malic enzymes (Mansouri *et al*, 2017; Montal *et al*, 2015), in Cluster 4 cells than in Cluster 3 (Fig 3I). This increased efflux of glutamine from the TCA cycle was also observed as greater enrichment of glutamine carbons into m+3 fructose 1,6-bisphosphate, and, to a lesser extent, phosphoenolpyruvate (Fig 3J), intermediates of gluconeogenesis. Combined, these data show that cells in Cluster 4 possessed a more oxidative phenotype that compensates for increased aerobic glycolysis with glutamine cataplerosis and explains the increased lactate production and more abundant TCA cycle metabolites in cells of Cluster 4, compared to Cluster 3 that had greater glucose and glutamine consumption and glucose oxidation (Fig 3K). Furthermore, these new insights into glucose and glutamine metabolism and the discovery that some cells produce more lactate despite lower glucose consumption support the new insights into high flux metabolic pathways from our pathway-centric ratio-based analyses.

### Differences in TCA cycle, lactate, and glutamine metabolism between Clusters 3 and 4 correlate with sensitivity to loss-of function

We further validated the outcomes of our pathway-centric metabolite ratio analysis of targeted metabolomic data (Fig 2) and functional studies (Fig 3) using *in silico* assessment of publicly available datasets. Specifically, we used the pan-cancer DEMETER (Tsherniak *et al*, 2017) and Project Score (Behan *et al*, 2019) loss-of-function screens, and the drug sensitivity PRISM (Corsello *et al*, 2020) and GDSC2 (Yang *et al*, 2012) datasets and selected for gene or drug targets within all KEGG metabolic pathways (Fig 4A). We then cross-referenced these large-scale cell line panels with cells that were members of Clusters 3 and 4 and focused our analyses to test the hypothesis that cells in Cluster 4 were more susceptible to depletion of genes associated with OXPHOS, glutamine and lactate metabolism compared to Cluster 3 (Fig 4B). In line with our hypothesis, Cluster 4 cells had greater sensitivity to genetic knockout of OXPHOS complexes I-V and glutaminase isoforms (Multiple unpaired t-tests, P<0.05; Fig 4C). We also observed a trend for greater sensitivity to deletion of succinate dehydrogenase (SDHD; Student t-test, P=0.06) and lactate dehydrogenase (LDHD; Student t-test, P=0.08) in cells in Cluster 4 (Appendix Fig S3A). We complemented our targeted assessment of these datasets by determining the top 20 metabolic targets from all KEGG pathways that possessed the greatest difference between Clusters 4 and 3. From this unbiased approach, we identified members of OXPHOS, glutamine metabolism, TCA cycle, and pyruvate/lactate pathways in this list (highlighted in yellow in Fig 4D, E). We also determined the top 20 list of the most different targets between Cluster 4 and 3 when we narrowed the coverage to just central carbon metabolism from all KEGG pathways (Appendix Fig S3B). Consistent with our other observations, the list of most sensitive targets again was enriched with enzymes from OXPHOS and TCA cycle pathways (Appendix Fig S3B). Finally, we developed a scoring system to consolidate the top 20 most sensitive pathways in Cluster 4 from all databases, and again found that the TCA cycle, OXPHOS, and glutamine metabolism were most vulnerable to genetic and drug targeting compared to Cluster 3 (Appendix Fig S3C). Combined, we have demonstrated that analyzing metabolomic data with a pathway-centric basis by using ratios identifies distinctive metabolic profiles that are evident in functional measures and loss-of-function screens.

**Figure 4:**
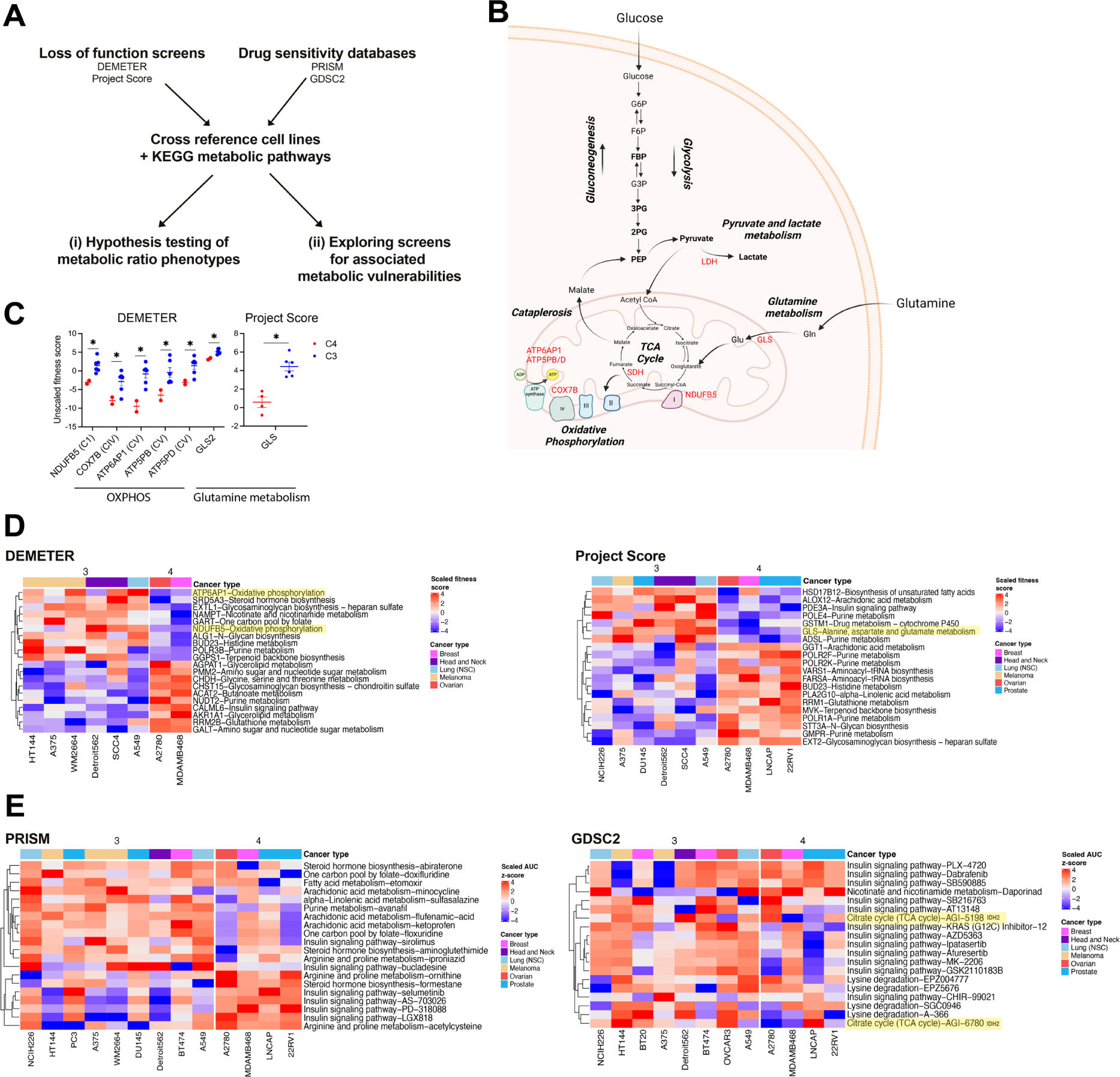
Validating pathway-centric metabolite ratio clusters and substrate dependencies using loss-of-function screens and drug sensitivity databases. A. Schematic of *in silico* analysis of loss-of-function and drug sensitivity screens against KEGG metabolic pathways and identified metabolic ratio phenotypes. B. Schematic illustrating the C4 phenotype of greater utilization of glucose and glutamine identified by cluster analyses using metabolite ratios. Genes named in red were hypothesized to have greater sensitivity to gene knockouts or inhibition in C4 compared to C3. C. Genes related to glucose and glutamine utilization phenotype of C4 with the greatest sensitivity to gene knockouts. (Multiple unpaired t-tests, *p<0.05, mean ± standard error of the mean, n=2-6 cell lines). D. Top 20 gene knockouts and associated KEGG pathways with the greatest fitness scores between C4 and C3 in DEMETER and Project Score databases. Genes and pathways matching C4 vulnerability model are highlighted in yellow. E. Top 20 drugs and associated KEGG pathways with the greatest differential AUC z-scores between C4 and C3 in PRISM and GDSC2 databases. Drugs and pathways matching C4 vulnerability model are highlighted in yellow.

## DISCUSSION

Cell metabolism is dynamic and differs depending on context. From the early observations of Warburg (Warburg, 1925; Warburg & Minami, 1923) and the Coris (Cori & Cori, 1925), it is now well accepted that cancer cells metabolize glucose differently from non-tumor tissue. These differences are not a consequence of rewiring or transformation of the cascades of biochemical reactions that form glycolysis and PPP but essentially from increased uptake of extracellular glucose driving increased flux and production of end products, including lactate (DeBerardinis & Chandel, 2020). Alongside these well-documented changes in glucose metabolism, there have been significant advances in the understanding of the interaction between metabolic pathways (Sung *et al*, 2023), outside the well-established convergence of glucose, glutamine and fatty acid metabolism in the TCA cycle. A major challenge remains how to infer mechanistic differences in metabolism based on metabolomics data alone, since the latter may not correlate with pathway activity nor flux. Conventional dimensional reduction, clustering, and over-presentation methodologies rely on coordinated changes to infer co-dependency or co-regulation (Amara *et al*, 2022; Huang & Wang, 2022), but the effectiveness of subsequent interpretations may be hinge on how the literature has delineated metabolite memberships (Mahajan *et al*, 2024), which are continuously honed over time.

Our first round of clustering, which was based on metabolite abundance of proliferating cells, formed groups that contained both epithelial and tumor-derived, and dismissed culturing conditions, tissue type and origin, and mutation status of oncogenic drivers as potential factors influencing cell metabolism. However, we failed to derive any metabolic signatures or hypotheses that could be evaluated with functional assessment. Consequently, we took a physiologically-based approach and transformed our metabolomic data into pathway-centric ratios and explored the relationships between reactant and product metabolites of central carbon and amino acid pathways. The approach draws upon thermodynamic principles in terms of reaction equilibrium and Gibbs energy (Park *et al*, 2016), and in practice, there is robust evidence for coordinated changes among proximal and hub metabolites (Martínez-Reyes & Chandel, 2020). Using metabolite ratios, we identified five clusters of cell lines, two of which (Cluster 3 and 4) we surmised to differ in TCA cycle activity based on the levels of TCA cycle metabolites relative to pyruvate. Indeed, the ensuing functional assay of glucose and glutamine metabolism verified pathway activity differences between Cluster 3 and 4, with Cluster 4’s elevated TCA cycle metabolites showing concordance with higher lactate production and the shift from glycolysis to an increased glutamine oxidation and cataplerosis. This work highlights the value of pathway-centric ratio-based data transformation in distinguishing metabolic pathway activity in a pan-cancer cell line panel and, therefore, supports the concept of a physiological-based approach to analyze metabolomic data.

Many tumors are reliant on glutamine as a critical carbon and nitrogen source, with glutaminase inhibition shown to be an effective therapy in several cancer types, such as triple-negative breast cancer (Gross *et al*, 2014), non-small cell lung cancer (van den Heuvel *et al*, 2012), and head and neck cancer (Wicker *et al*, 2021). Our findings showed that Cluster 4 cells had higher OXPHOS levels yet consumed less glucose and glutamine; these cells appear more carbon efficient. The relatively more plentiful TCA cycle metabolites may sustain higher TCA cycle fluxes, which are tightly coupled to OXPHOS, and buffer critical anabolic and signaling functions (Martínez-Reyes & Chandel, 2020). Glutamine’s proximity means oxidizing glutamine repletes TCA cycle metabolites more directly than glucose (Quek *et al*, 2022), but the displacement of glucose oxidation further entrenched the aerobic glycolysis phenotype as seen in Cluster 4 cells. Additionally, glutamine cataplerosis and the contribution of PEP carboxykinase to gluconeogenesis, converting oxaloacetate to PEP, augment the supply of biomass precursors, which have been documented in several cancer types, including liver (Liu *et al*, 2018) and lung (Vincent *et al*, 2015). For example, glutamine-derived lactate production via glutaminolysis in glioblastoma cells helps produce NADPH and support fatty acid synthesis (DeBerardinis *et al*, 2007). Overall, we speculate that the increased glutamine utilization among Cluster 4 cells may confer greater fitness to support proliferation and survival.

Excitingly, our assertion of elevated OXPHOS and enhanced glutamine utilization in Cluster 4 matched data mining results derived from drug sensitivity (Corsello *et al*., 2020; Yang *et al*., 2012) and loss-of-function screens (Behan *et al*., 2019; Tsherniak *et al*., 2017). Among the top 20 loss-of-function screens, Cluster 4 cells were most sensitive to targeting the TCA cycle, OXPHOS, glycolysis, and glutamate metabolism pathways, compared to Cluster 3 cells. Namely, we found Cluster 4 cells significantly more sensitive to glutaminase gene knockout and inhibitors. These findings highlight the potential of a physiological pathway-centric approach to translating metabolite signatures into effective strategies for identifying druggable vulnerabilities in glucose and glutamine metabolism.

A key limitation of our approach is coverage. Our targeted LC-MS method covers central carbon metabolism, and thus we have focused on the TCA cycle as it is a convergent point for glucose and glutamine utilization; however, whether these relationships are consistent for other TCA cycle substrates, such as branched-chain amino acids and fatty acids (Neinast *et al*, 2019; Schoors *et al*, 2015) remains to be determined. Another major lesson from our approach was that the most insightful outcome of our clustering analysis came from using a hub metabolite (e.g., pyruvate) as the denominator rather than a starting substrate (i.e., glucose, glutamine, serine). Since we only included one hub metabolite in our pathway cluster analysis, it is conceivable that investigating other metabolic hubs not covered in our targeted approach, such as NAD^+^ (Benedetti *et al*., 2023), could identify other distinctive signatures and targetable vulnerabilities. Perhaps the abundance of hub metabolites, where multiple pathways converge, are less prone to isolated variation or are tightly regulated and thus more effective at distinguishing metabolic changes in a ratio approach.

A major motivation for the current study was to identify common metabolic signatures that arise from various genomic bases that can form the foundation for a simpler therapeutic approach across cancer types. The outcomes of this study have, in part, provided evidence that this may be achievable. Our results highlight the potential of using physiologically-based, pathway-centric metabolite ratios to gain insights into the convergent or recurrent pathway mechanisms within subsets of diverse cancer types and identify targetable vulnerabilities. Combined with existing large-scale metabolomic datasets, our approach may lay the foundation to accelerate the ongoing efforts to profile cancer metabolism for future therapeutic advances and repurposing metabolic targeting-based therapeutics in a pan-cancer setting.

## MATERIALS AND METHODS

### Reagents and Tools Table

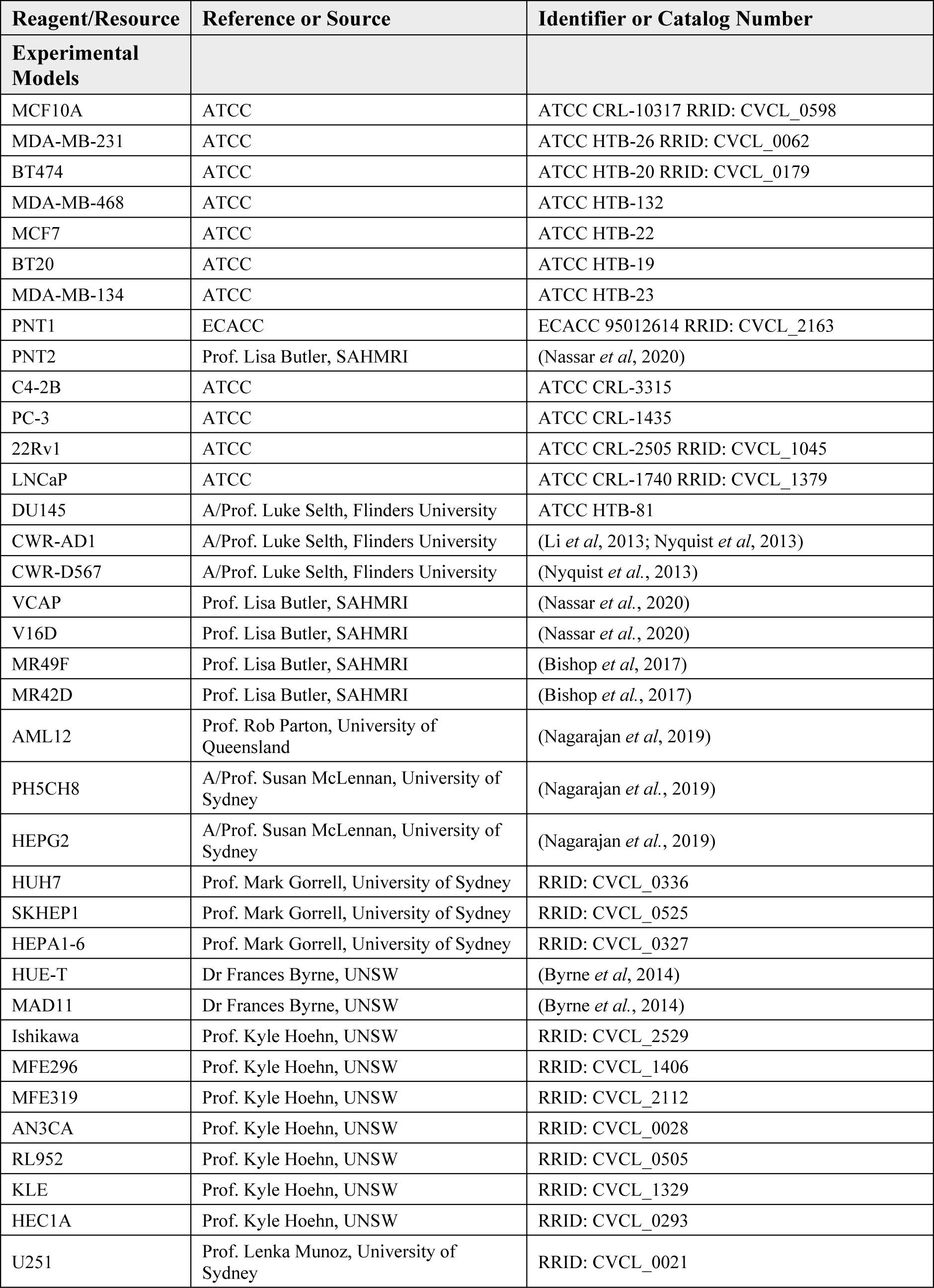
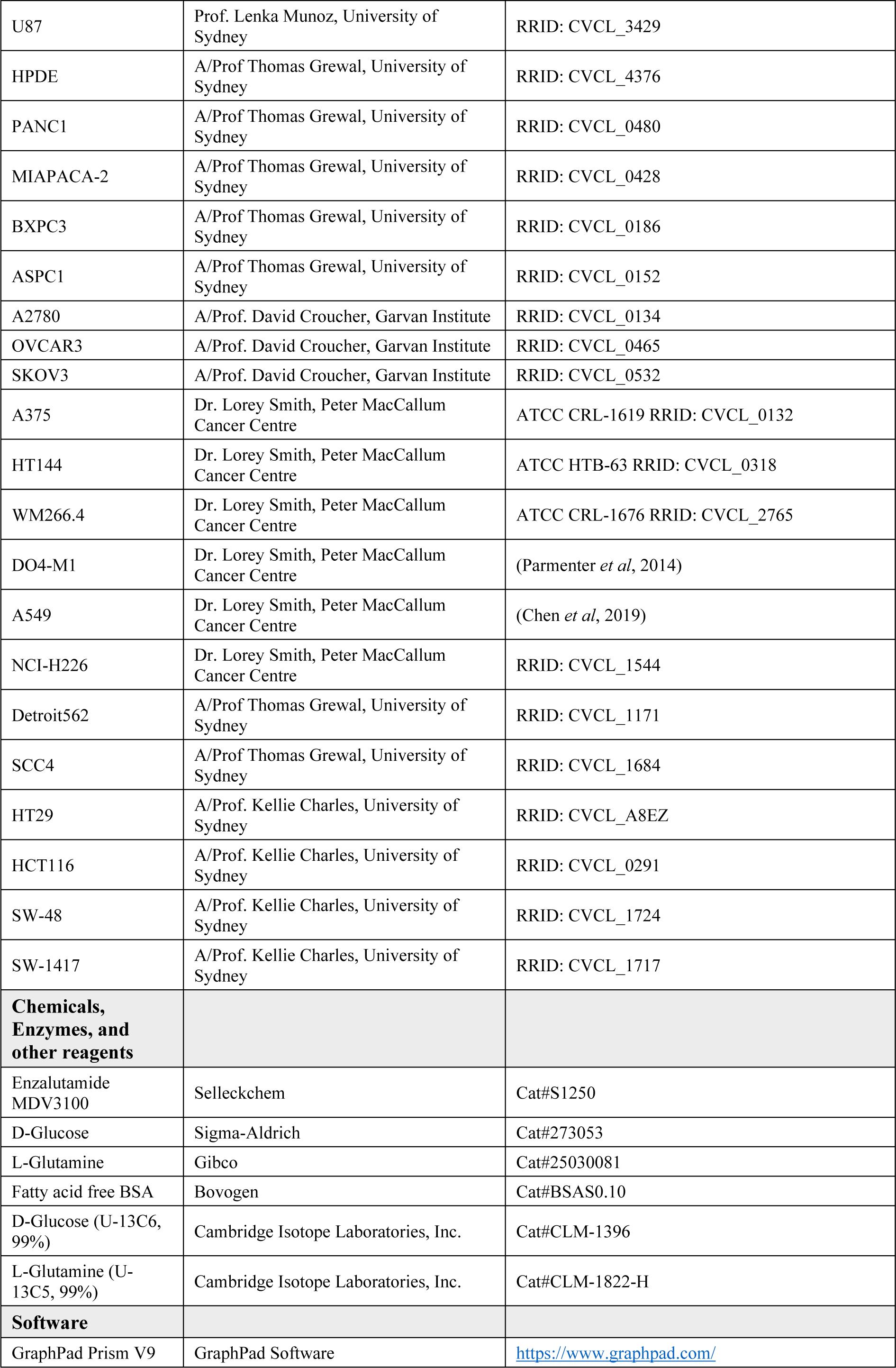
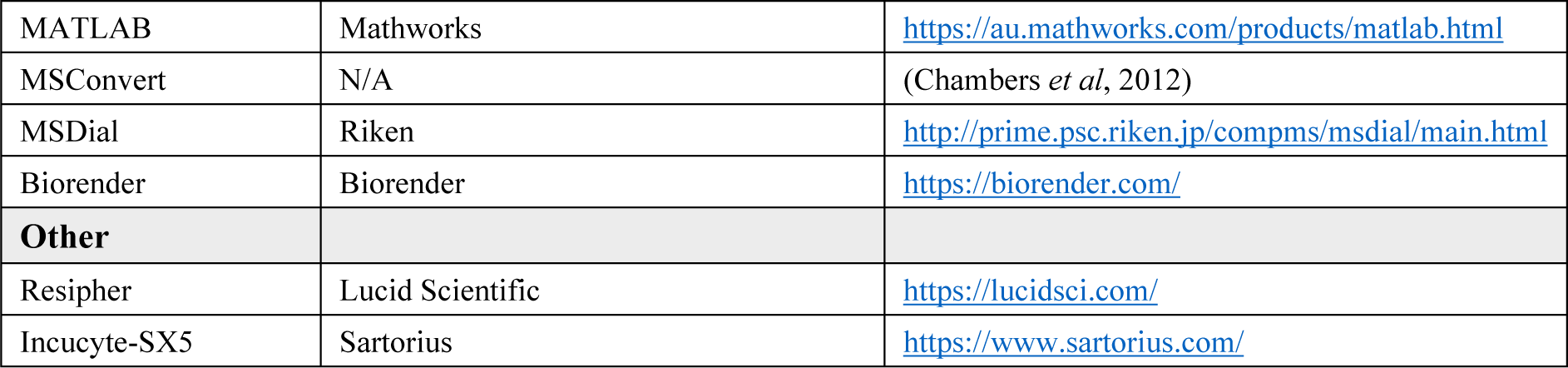

### Cell lines and culture conditions

A total of 57 adherent cell lines (8 normal and 49 tumor) spanning 11 cancer types were used in this study. Cell lines obtained from vendors or that were kindly provided by academic labs, overlapped with major cell line resources such as the Cancer Cell Line Encyclopedia (CCLE) and National Cancer Institute 60 panel (NCI60) (Appendix table 1). Mutational status of common oncogenes was characterized by cross-referencing the Cellosaurus database, COSMIC database, and Depmap portal. Culturing media were supplemented with 10% fetal bovine serum (Cytiva Hyclone) and 1% penicillin/streptomycin (Gibco) unless stated otherwise (Appendix table 1). Cells were incubated at 37°C and 5% CO_2_. The full panel of cells were generated across 6 batches over a period of 3 years due to COVID restrictions and included overlapping cell lines to correct for batch effects.

### Metabolomics experiments

Cells were seeded in triplicate in 6-well plates at a density of 5×10^5^ cells/well in 2 mL of media. After 24 hours, the media was removed, and cells were washed once with 2 mL ice-cold 0.9% w/v NaCl. Cells were then scraped with 300 mL of extraction buffer, EB, (1:1 LC/MS methanol:water (Optima) + 0.1x internal standards comprised of non-endogenous polar metabolites (2-morpholinoethanesulfonic acid, D-camphor-10-sulfonic acid, and deuterated thymidine) and transferred to a 1.5 mL microcentrifuge tube. A further 300 mL of EB was added to the well and combined in the tube. 600 mL Chloroform (Honeywell) was added before vortexing and incubating on ice for 10 minutes. Tubes were vortexed briefly and centrifuged at 15,000 x *g* for 10 minutes at 4°C. The aqueous layer was collected and dried without heat, using a Savant SpeedVac (Thermo Fisher). Dried samples were resuspended in 40 mL Amide buffer A (20 mM ammonium acetate, 20 mM ammonium hydroxide, 95:5 HPLC H_2_O: Acetonitrile (v/v)) and vortexed and centrifuged at 15,000 x *g* for 5 minutes at 4°C. 20 mL of supernatant was transferred to HPLC vials containing 20 mL acetonitrile for LC-MS analysis of amino acids and glutamine metabolites. The remaining 20 mL of resuspended sample was transferred to HPLC vials containing 20 mL LC-MS H_2_O for LCMS analysis of glycolytic, pentose phosphate pathway, and TCA cycle metabolites. Amino acids and glutamine metabolites were measured using the Vanquish-TSQ Altis (Thermo) LC-MS/MS system. Analyte separation was achieved using a Poroshell 120 HILIC-Z Column (2.1×150 mm, 2.7 mm) (Agilent) at ambient temperature. The pair of buffers used were Amide buffer A and 100% acetonitrile (Buffer B), flowed at 200 mL/min; injection volume of 5 mL. Glycolytic, PPP and TCA cycle metabolites were measured using 1260 Infinity (Agilent)-QTRAP6500+ (AB Sciex) LC-MS/MS system. Analyte separation was achieved using a Synergi 2.5 mm Hydro-RP 100A LC Column (100×2 mm) at ambient temperature. The pair of buffers used were 95:5 (v/v) water:acetonitrile containing 10 mM tributylamine and 15 mM acetic acid (Buffer A) and 100% acetonitrile (Buffer B), flowed at 200 mL/min; injection volume of 5 mL. Raw data from both LC-MS/MS systems were extracted using MSConvert (Chambers *et al*., 2012) and in-house MATLAB scripts. Concentrations of metabolites were calculated against a standard curve of polar and amino acid metabolite standards similarly extracted as above. Log10 normalization was performed on the metabolite concentration data.

### Metabolomics batch correction

Cell line samples were quantified over 6 batches during the experiment including overlapping cell lines across each batch. To eliminate potential batch effects, we applied the normalization method: Removing Unwanted Variation-III (Molania *et al*, 2019). We set k, the number of unwanted factors, to 9. The cell lines that were measured across multiple runs were used as replicates for the batch correction algorithm and to assess the quality of the batch correction output.

### Metabolomics clustering analysis

Clustering was performed on the batch-corrected matrix using K-means algorithms with “1 - Pearson correlation” as the distance matrix (Gu *et al*, 2016) implemented in the ComplexHeatmap package (k = 5). We visualized the results as a heatmap using the ComplexHeatmap package. The clustering procedure was repeated 1000 times and the consensus was taken to ensure a more robust clustering result. The clustering strategy was applied to both the original batch-corrected matrix and the ratio-transformed matrix. We visually assess whether the obtained clustering was not influenced by potential confounding factors, such as culturing conditions and common oncogenic gene mutations and tissue type. This is achieved by adding a color-coded legend on culturing conditions and mutational information to the clustering heatmap and demonstrate that no patterns exist.

The original batch-corrected matrix was subset to targeted metabolites from several key metabolic pathways including the TCA cycle, pentose phosphate pathway, glycolysis, amino acid metabolism, and glutamine metabolism, followed by clustering analysis. For the ratio-transformed matrix, batch-corrected abundance ratios were calculated between a precursor metabolite for each pathway (used as the denominator) and the remaining pathway metabolites. The corresponding precursor (denominator) metabolites were pyruvate for the TCA cycle, glucose for PPP and glycolysis, and glutamine for the glutamine metabolism pathway. For amino acid pathways, serine, proline, and methionine were used as the precursor metabolites. The clustering structure determined from the processes above were also used to visualize the oncogene mutation status, tissue type, cancer type, tissue origin, and culturing media type of the cell lines using a heatmap and inspect the relationship of these variables with the determined clusters.

### U-^13^C stable isotope tracing

Cells were seeded in triplicate in 6-well plates at a density of 5×10^5^ cells/well in 2 mL of media. After 24 hours wells were washed with warm PBS and media replaced with 600 mL of DMEM no glucose, no glutamine media supplemented with 2% (wt./vol) FA-free BSA, and 5 mM glucose and 1 mM glutamine, replaced with their respective U-^13^C forms. Cells were incubated in U-^13^C containing medium for 6 hours. Samples were extracted and measured using the Vanquish-TSQ Altis and Agilent-QTRAP6500+.

### Extracellular substrate experiments

Cells were seeded in triplicate in 6-well plates at a density of 5×10^5^ cells/well in 2 mL of media. After 24 hours wells were washed with warm PBS and media replaced with 1 mL of DMEM no glucose, no glutamine media supplemented with 5 mM glucose, 1 mM glutamine and 150 mM palmitate. 100 mL of extracellular media were collected from wells at 3, 6, 12, 24 hour timepoints. Media samples were centrifuged at 16,000 x *g* for 5 minutes at 4°C and the supernatant collected for subsequent extraction and LC-MS analysis.

To extract media samples for LC-MS, 20 mL of supernatant media was first diluted with 80 mL water and vortexed, and then 10mL of the diluted media was transferred to 90 mL of extraction buffer containing 1:1 (v/v) acetonitrile and methanol + 1x internal standards (non-endogenous standards) at −30°C. The mixture was centrifuged at 12,000 x *g* for 5 minutes at 4°C and transferred into HPLC vials for LC-MS analysis measured using the Vanquish-TSQ Altis.

### Oxygen consumption rate

Cells were seeded in 96-well plates (Nunc) in 100 mL basal medium. Oxygen consumption rates were continuously measured using Resipher (Lucid Scientific) at 37°C, 10% CO_2_ as per manufacturer instructions over 48 hours, starting 24 hours post-seeding. Media was replenished at 24 hours. Cell-free wells contained 200 mL of PBS to avoid evaporation. To account for differences in cell growth over 48 hours, parallel plates were similarly cultured, and media replenished at 24 hours for cell confluency measured using IncuCyte-SX5 (Sartorius).

### Analysis of drug and CRISPR databases

Drug sensitivity area under the curve (AUC) data were downloaded from the PRISM (Corsello *et al*., 2020) and GDSC2 (Release 8.4, July 2022) (Yang *et al*., 2012) databases. Loss-of-function fitness score data were downloaded from the DEMETER (Tsherniak *et al*., 2017) and Project Score (July 2021) (Behan *et al*., 2019) databases. First, cell lines were filtered by overlapping cell lines within our panel, and then inhibitor or loss-of-function gene targets were filtered by KEGG metabolic pathway genes. We then performed a differential expression analysis on the drug response of the cell lines belonging to clusters of interest, specifically the response of Cluster 3 cell lines versus Cluster 4 cell lines. The top 20 drugs or gene targets with the greatest differential response between Clusters 3 and 4 was identified.

### Statistical analysis for *in vitro* and database analysis

For all ^13^C-tracing, extracellular, and oxygen consumption experiments, at least 3 technical replicates and 3 independent biological replicates were used for each sample group. Descriptive data summary in Figures 3 and 4 were presented as mean ± standard error of the mean (SEM), mean ± standard deviation (SD), or mean ± min to max values, as indicated in each of the figure legends. We determine statistically significant differences between cell clusters 3 and 4 in ^13^C-tracing, extracellular, and oxygen consumption experiments and loss-of-function analysis by performing unpaired Student’s t-test. We assessed the differences between clusters 3 and 4 of oxidative phosphorylation gene loss-of-function by multiple unpaired t-tests (implemented in Prism GraphPad V9). Statistical significance is indicated in all figures by the following annotations *p <0.05 and not statistically different otherwise. The R software and packages was used for clustering methods and heatmaps as reported in previous sections. Schematic diagrams were created with Biorender.com.

---

**Appendix Figure S1:**
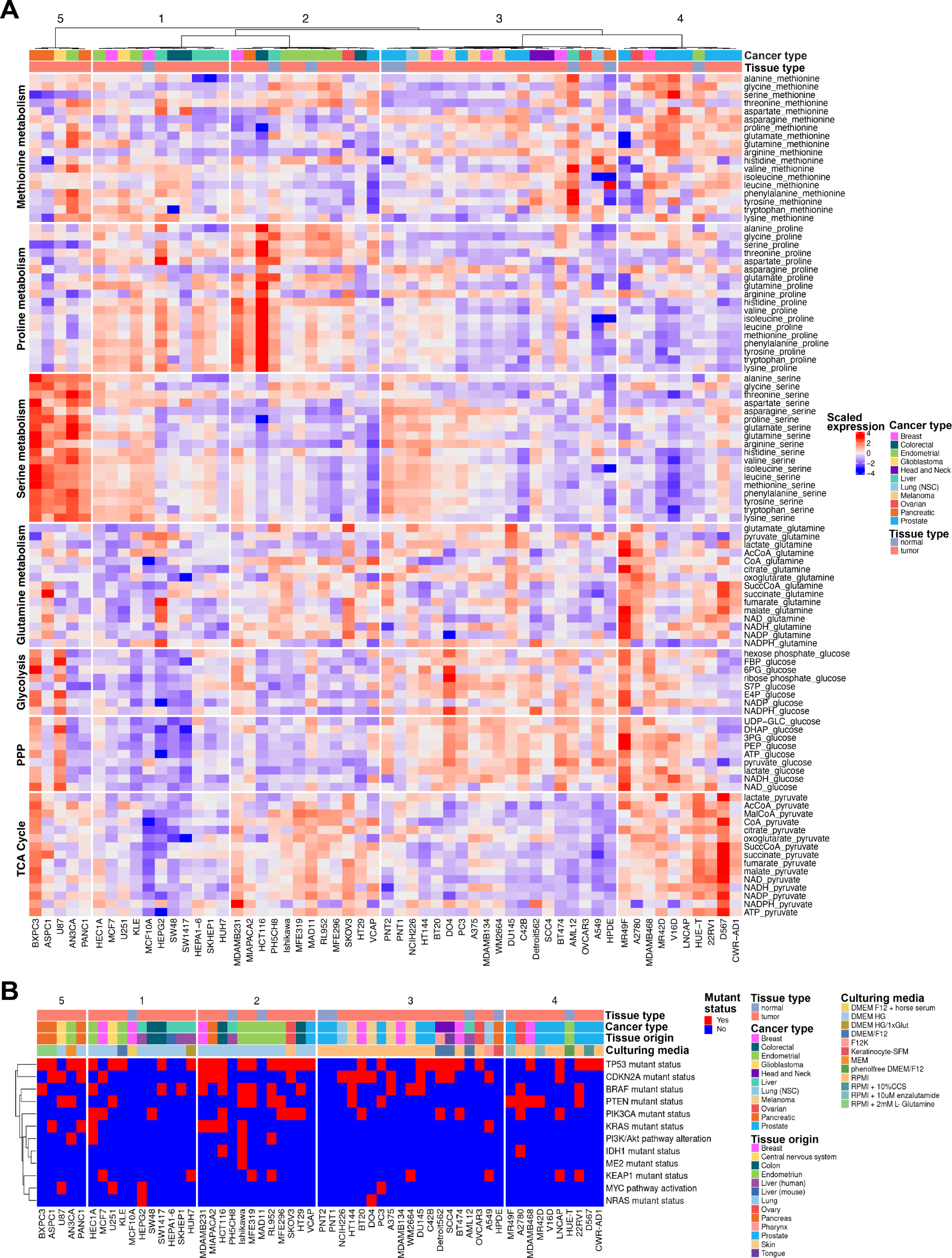
Pathway-centric metabolite ratios of all targeted metabolic pathways. A. Heatmap of scaled metabolite ratios by pathways covered in the targeted metabolomics approach. Ratios calculated for all metabolites of a specific pathway against a precursor metabolite. B. Clusters of cell lines from (A) appended with color coded legends for mutant status of common oncogenic drivers, culturing media conditions and tissue origins.

**Appendix Figure S2:**
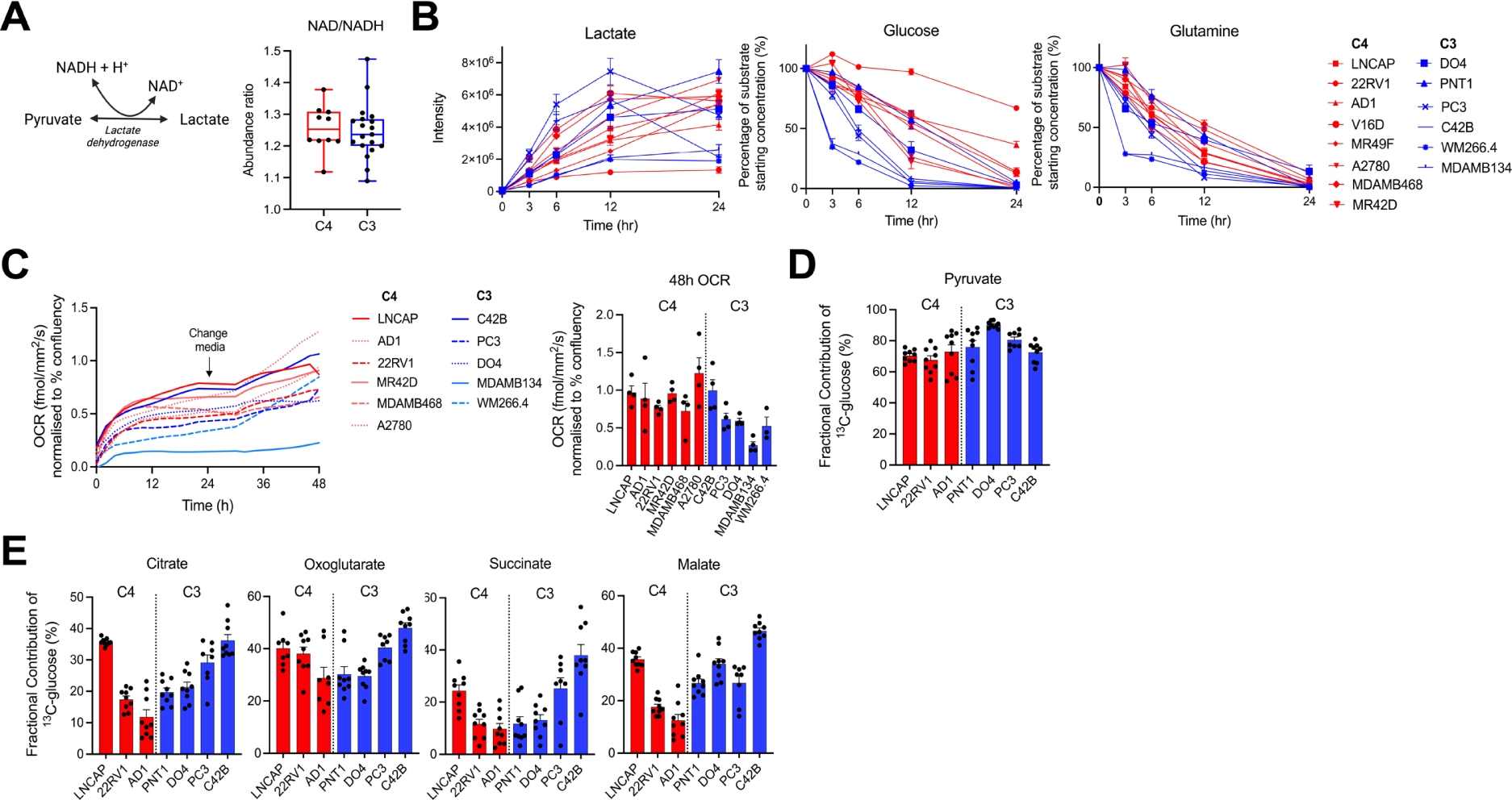
Substrate preferences for cell line clusters identified by TCA cycle ratios. A. Schematic of the lactate dehydrogenase catalyzed reaction. NAD+ to NADH ratio in cells of Clusters 4 and 3 (Unpaired student t-test, ns p>0.05, error min to max values). B. Lactate production and glucose and glutamine consumption over 24 hours (C4 n=8, C3 n-6, 3 biological replicates and 3 technical replicates per cell line, mean ± standard error of the mean). C. Oxygen consumption rates (OCR) measured over 48 hours, normalized to % confluency. Media changed at 24 hours (left). OCR measurement for cell lines at 48 hours (right) (C4 n=6, C3 n-5, 3 biological replicates and 4 technical replicates per cell line). D. Fractional contribution of [U-^13^C]-glucose to intracellular pyruvate (C4 n=3, C3 n=4, 3 biological replicates and 3 technical replicates per cell line, mean ± standard error of the mean). E. Fractional contribution of [U-^13^C]-glucose to TCA metabolites citrate, oxoglutarate, succinate and malate (C4 n=3, C3 n=4, 3 biological replicates and 3 technical replicates per cell line, mean ± standard error of the mean).

**Appendix Figure S3:**
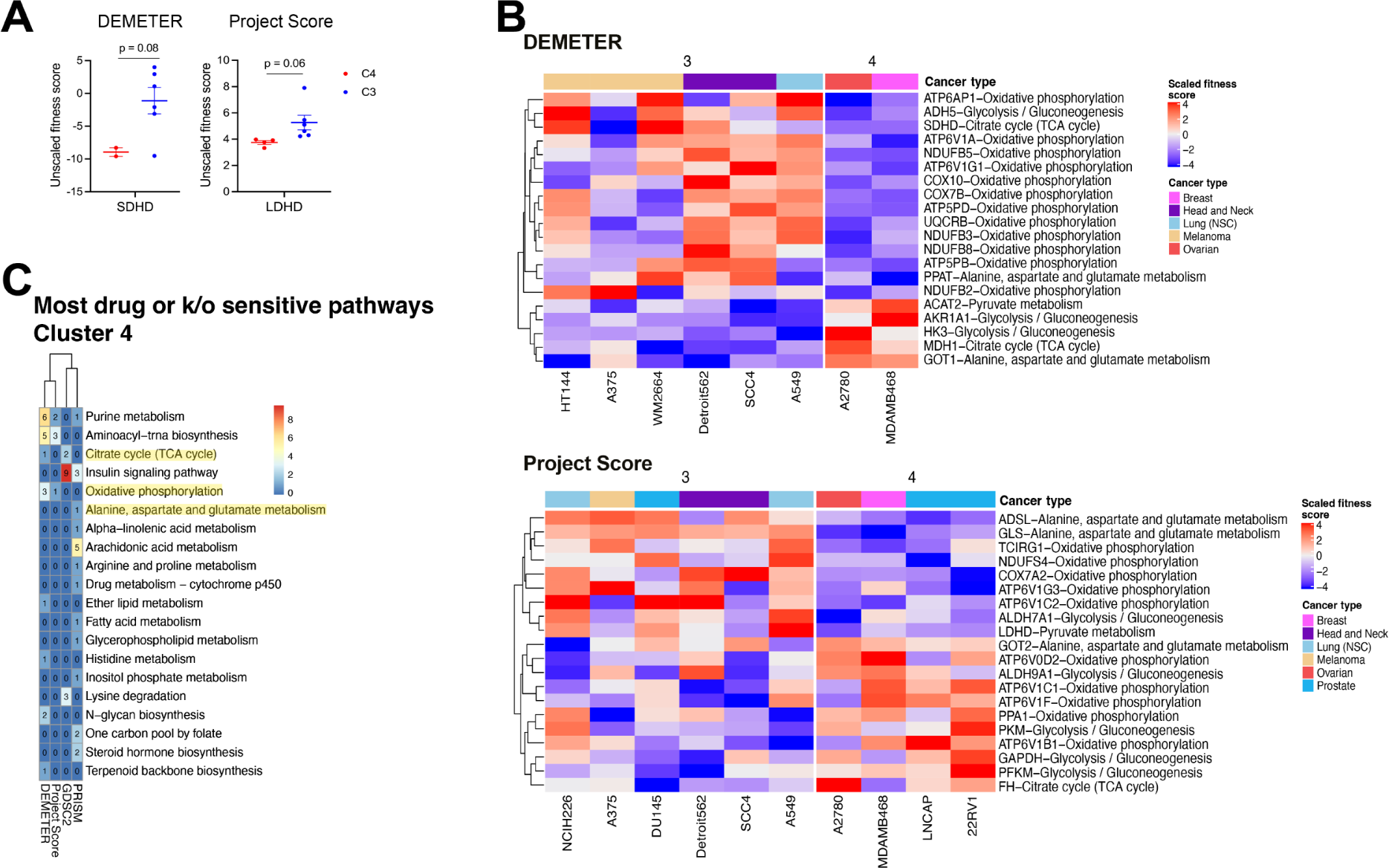
Loss of function and drug database screen validation of pathway-centric metabolite ratio clusters. A. Genes related to the glucose and glutamine utilization phenotype of C4 with trends of greater sensitivity to gene knockouts. (Unpaired student t-test, ns p>0.05, mean ± standard error of the mean, n=2-6). B. Top 20 gene knockouts and associated KEGG pathways with the greatest fitness scores between C4 and C3 in DEMETER and Project Score databases from glycolysis, TCA cycle, pyruvate metabolism, glycolysis/gluconeogenesis, and alanine, aspartate, and glutamate metabolism pathways. C. Metabolic pathways associated with the top 20 drug and gene knockout sensitives for Cluster 4 for each database. Drugs and pathways matching C4 vulnerability model (yellow) are highlighted.

## Acknowledgements

N.T.S. is supported by the Australian Rotary Health/Rotary Club of Blacktown City ‘Mel Grey’ PhD scholarship. A.J.H. is supported by a Robinson Fellowship. This work was supported by funding from The University of Sydney. We thank the Sydney Mass Spectrometry facility for access to LC-MS instruments; and all collaborators who kindly donated cell lines.

## Author contributions

### Nancy T Santiappillai

Project administration, formal analysis, investigation, validation, visualization, writing – original draft, writing -review & editing. **Yue Cao:** Formal analysis, methodology, software, validation, visualization, writing -review & editing. **Mariam F Hakeem-Sanni:** Investigation. **Jean Yang:** Methodology, supervision, writing -review & editing. **Lake-Ee Quek:** Conceptualization, methodology, project administration, software, supervision, writing – review & editing. **Andrew J Hoy:** Conceptualization, funding acquisition, project administration, resources, supervision, writing – review & editing.

## REFERENCES

Altea-Manzano P, Cuadros AM, Broadfield LA, Fendt SM (2020) Nutrient metabolism and cancer in the in vivo context: a metabolic game of give and take. EMBO Rep 21: e50635

Amara A, Frainay C, Jourdan F, Naake T, Neumann S, Novoa-del-Toro EM, Salek RM, Salzer L, Scharfenberg S, Witting M (2022) Networks and Graphs Discovery in Metabolomics Data Analysis and Interpretation. Frontiers in Molecular Biosciences 9

Behan FM, Iorio F, Picco G, Gonçalves E, Beaver CM, Migliardi G, Santos R, Rao Y, Sassi F, Pinnelli M et al (2019) Prioritization of cancer therapeutic targets using CRISPR–Cas9 screens. Nature 568: 511–516

Benedetti E, Liu EM, Tang C, Kuo F, Buyukozkan M, Park T, Park J, Correa F, Hakimi AA, Intlekofer AM et al (2023) A multimodal atlas of tumour metabolism reveals the architecture of gene–metabolite covariation. Nat Metab 5: 1029–1044

Bishop JL, Thaper D, Vahid S, Davies A, Ketola K, Kuruma H, Jama R, Nip KM, Angeles A, Johnson F et al (2017) The Master Neural Transcription Factor BRN2 Is an Androgen Receptor–Suppressed Driver of Neuroendocrine Differentiation in Prostate Cancer. Cancer Discov 7: 54–71

Byrne FL, Poon IKH, Modesitt SC, Tomsig JL, Chow JDY, Healy ME, Baker WD, Atkins KA, Lancaster JM, Marchion DC et al (2014) Metabolic Vulnerabilities in Endometrial Cancer. Cancer Res 74: 5832–5845

Cairns RA, Harris IS, Mak TW (2011) Regulation of cancer cell metabolism. Nat Rev Cancer 11: 85–95

Chambers MC, Maclean B, Burke R, Amodei D, Ruderman DL, Neumann S, Gatto L, Fischer B, Pratt B, Egertson J et al (2012) A cross-platform toolkit for mass spectrometry and proteomics. Nat Biotechnol 30: 918–920

Chen P-H, Cai L, Huffman K, Yang C, Kim J, Faubert B, Boroughs L, Ko B, Sudderth J, McMillan EA et al (2019) Metabolic Diversity in Human Non-Small Cell Lung Cancer Cells. Mol Cell 76: 838–851.e835

Cherkaoui S, Durot S, Bradley J, Critchlow S, Dubuis S, Masiero MM, Wegmann R, Snijder B, Othman A, Bendtsen C et al (2022) A functional analysis of 180 cancer cell lines reveals conserved intrinsic metabolic programs. Mol Syst Biol 18

Cori CF, Cori GT (1925) the carbohydrate metabolism of tumors: ii. Changes in the sugar, lactic acid, and co2-combining power of blood passing through a tumor. J Biol Chem 65: 397–405

Corsello SM, Nagari RT, Spangler RD, Rossen J, Kocak M, Bryan JG, Humeidi R, Peck D, Wu X, Tang AA et al (2020) Discovering the anti-cancer potential of non-oncology drugs by systematic viability profiling. Nat Cancer 1: 235–248

DeBerardinis RJ, Chandel NS (2020) We need to talk about the Warburg effect. Nat Metab 2: 127–129

DeBerardinis RJ, Mancuso A, Daikhin E, Nissim I, Yudkoff M, Wehrli S, Thompson CB (2007) Beyond aerobic glycolysis: transformed cells can engage in glutamine metabolism that exceeds the requirement for protein and nucleotide synthesis. Proc Natl Acad Sci U S A 104: 19345–19350

Fendt SM, Frezza C, Erez A (2020) Targeting Metabolic Plasticity and Flexibility Dynamics for Cancer Therapy. Cancer Discov 10: 1797–1807

Gross MI, Demo SD, Dennison JB, Chen L, Chernov-Rogan T, Goyal B, Janes JR, Laidig GJ, Lewis ER, Li J et al (2014) Antitumor Activity of the Glutaminase Inhibitor CB-839 in Triple-Negative Breast Cancer. Mol Cancer Ther 13: 890–901

Gu Z, Eils R, Schlesner M (2016) Complex heatmaps reveal patterns and correlations in multidimensional genomic data. Bioinformatics 32: 2847–2849

Hensley CT, Faubert B, Yuan Q, Lev-Cohain N, Jin E, Kim J, Jiang L, Ko B, Skelton R, Loudat L et al (2016) Metabolic Heterogeneity in Human Lung Tumors. Cell 164: 681–694

Huang Z, Wang C (2022) A Review on Differential Abundance Analysis Methods for Mass Spectrometry-Based Metabolomic Data. Metabolites 12

Jain M, Nilsson R, Sharma S, Madhusudhan N, Kitami T, Souza AL, Kafri R, Kirschner MW, Clish CB, Mootha VK (2012) Metabolite profiling identifies a key role for glycine in rapid cancer cell proliferation. Science 336: 1040–1044

Jia S, Liu Z, Zhang S, Liu P, Zhang L, Lee SH, Zhang J, Signoretti S, Loda M, Roberts TM et al (2008) Essential roles of PI(3)K–p110β in cell growth, metabolism and tumorigenesis. Nature 454: 776–779

Jones RG, Thompson CB (2009) Tumor suppressors and cell metabolism: a recipe for cancer growth. Genes Dev 23: 537–548

Kamphorst JJ, Nofal M, Commisso C, Hackett SR, Lu W, Grabocka E, Vander Heiden MG, Miller G, Drebin JA, Bar-Sagi D et al (2015) Human pancreatic cancer tumors are nutrient poor and tumor cells actively scavenge extracellular protein. Cancer Res 75: 544–553

Li H, Ning S, Ghandi M, Kryukov GV, Gopal S, Deik A, Souza A, Pierce K, Keskula P, Hernandez D et al (2019) The landscape of cancer cell line metabolism. Nat Med 25: 850–860

Li Y, Chan SC, Brand LJ, Hwang TH, Silverstein KA, Dehm SM (2013) Androgen receptor splice variants mediate enzalutamide resistance in castration-resistant prostate cancer cell lines. Cancer Res 73: 483–489

Liu M-X, Jin L, Sun S-J, Liu P, Feng X, Cheng Z-L, Liu W-R, Guan K-L, Shi Y-H, Yuan H-X et al (2018) Metabolic reprogramming by PCK1 promotes TCA cataplerosis, oxidative stress and apoptosis in liver cancer cells and suppresses hepatocellular carcinoma. Oncogene 37: 1637–1653

Luengo A, Li Z, Gui DY, Sullivan LB, Zagorulya M, Do BT, Ferreira R, Naamati A, Ali A, Lewis CA et al (2021) Increased demand for NAD+ relative to ATP drives aerobic glycolysis. Mol Cell 81: 691–707.e696

Mahajan P, Fiehn O, Barupal D (2024) IDSL.GOA: gene ontology analysis for interpreting metabolomic datasets. Sci Rep 14: 1299

Mansouri S, Shahriari A, Kalantar H, Moini Zanjani T, Haghi Karamallah M (2017) Role of malate dehydrogenase in facilitating lactate dehydrogenase to support the glycolysis pathway in tumors. Biomed Rep 6: 463–467

Martínez-Reyes I, Chandel NS (2020) Mitochondrial TCA cycle metabolites control physiology and disease. Nat Commun 11

Molania R, Gagnon-Bartsch JA, Dobrovic A, Speed TP (2019) A new normalization for Nanostring nCounter gene expression data. Nucleic Acids Res 47: 6073–6083

Montal ED, Dewi R, Bhalla K, Ou L, Hwang BJ, Ropell AE, Gordon C, Liu WJ, DeBerardinis RJ, Sudderth J et al (2015) PEPCK Coordinates the Regulation of Central Carbon Metabolism to Promote Cancer Cell Growth. Mol Cell 60: 571–583

Mullen NJ, Singh PK (2023) Nucleotide metabolism: a pan-cancer metabolic dependency. Nat Rev Cancer 23: 275–294

Nagarajan SR, Paul-Heng M, Krycer JR, Fazakerley DJ, Sharland AF, Hoy AJ (2019) Lipid and glucose metabolism in hepatocyte cell lines and primary mouse hepatocytes: a comprehensive resource for in vitro studies of hepatic metabolism. Am J Physiol Endocrinol Metab 316: E578–E589

Nassar ZD, Mah CY, Dehairs J, Burvenich IJ, Irani S, Centenera MM, Helm M, Shrestha RK, Moldovan M, Don AS et al (2020) Human DECR1 is an androgen-repressed survival factor that regulates PUFA oxidation to protect prostate tumor cells from ferroptosis. Elife 9

Neinast MD, Jang C, Hui S, Murashige DS, Chu Q, Morscher RJ, Li X, Zhan L, White E, Anthony TG et al (2019) Quantitative Analysis of the Whole-Body Metabolic Fate of Branched-Chain Amino Acids. Cell Metab 29: 417–429.e414

Nyquist MD, Li Y, Hwang TH, Manlove LS, Vessella RL, Silverstein KAT, Voytas DF, Dehm SM (2013) TALEN-engineered AR gene rearrangements reveal endocrine uncoupling of androgen receptor in prostate cancer. Proc Natl Acad Sci U S A 110: 17492–17497

Oermann EK, Wu J, Guan KL, Xiong Y (2012) Alterations of metabolic genes and metabolites in cancer. Semin Cell Dev Biol 23: 370–380

Ortmayr K, Dubuis S, Zampieri M (2019) Metabolic profiling of cancer cells reveals genome-wide crosstalk between transcriptional regulators and metabolism. Nat Commun 10

Park JO, Rubin SA, Xu YF, Amador-Noguez D, Fan J, Shlomi T, Rabinowitz JD (2016) Metabolite concentrations, fluxes and free energies imply efficient enzyme usage. Nat Chem Biol 12: 482–489

Parmenter TJ, Kleinschmidt M, Kinross KM, Bond ST, Li J, Kaadige MR, Rao A, Sheppard KE, Hugo W, Pupo GM et al (2014) Response of BRAF-mutant melanoma to BRAF inhibition is mediated by a network of transcriptional regulators of glycolysis. Cancer Discov 4: 423–433

Pavlova NN, Zhu J, Thompson CB (2022) The hallmarks of cancer metabolism: Still emerging. Cell Metab 34: 355–377

Quek L-E, van Geldermalsen M, Guan YF, Wahi K, Mayoh C, Balaban S, Pang A, Wang Q, Cowley MJ, Brown KK et al (2022) Glutamine addiction promotes glucose oxidation in triple-negative breast cancer. Oncogene 41: 4066–4078

Schoors S, Bruning U, Missiaen R, Queiroz KC, Borgers G, Elia I, Zecchin A, Cantelmo AR, Christen S, Goveia J et al (2015) Fatty acid carbon is essential for dNTP synthesis in endothelial cells. Nature 520: 192–197

Shorthouse D, Bradley J, Critchlow SE, Bendtsen C, Hall BA (2022) Heterogeneity of the cancer cell line metabolic landscape. Mol Syst Biol 18

Smith B, Schafer XL, Ambeskovic A, Spencer CM, Land H, Munger J (2016) Addiction to Coupling of the Warburg Effect with Glutamine Catabolism in Cancer Cells. Cell Rep 17: 821–836

Spinelli JB, Haigis MC (2018) The multifaceted contributions of mitochondria to cellular metabolism. Nat Cell Biol 20: 745–754

Sung Y, Yu YC, Han JM (2023) Nutrient sensors and their crosstalk. Exp Mol Med 55: 1076–1089

Tsherniak A, Vazquez F, Montgomery PG, Weir BA, Kryukov G, Cowley GS, Gill S, Harrington WF, Pantel S, Krill-Burger JM et al (2017) Defining a Cancer Dependency Map. Cell 170: 564–576.e516

van den Heuvel AP, Jing J, Wooster RF, Bachman KE (2012) Analysis of glutamine dependency in non-small cell lung cancer: GLS1 splice variant GAC is essential for cancer cell growth. Cancer Biol Ther 13: 1185–1194

Vincent Emma E, Sergushichev A, Griss T, Gingras M-C, Samborska B, Ntimbane T, Coelho Paula P, Blagih J, Raissi Thomas C, Choinière L et al (2015) Mitochondrial Phosphoenolpyruvate Carboxykinase Regulates Metabolic Adaptation and Enables Glucose-Independent Tumor Growth. Mol Cell 60: 195–207

Vousden KH, Ryan KM (2009) p53 and metabolism. Nat Rev Cancer 9: 691–700

Wang P, Ma J, Li W, Wang Q, Xiao Y, Jiang Y, Gu X, Wu Y, Dong S, Guo H et al (2023) Profiling the metabolome of uterine fluid for early detection of ovarian cancer. Cell Rep Med: 101061

Warburg O (1925) über den Stoffwechsel der Carcinomzelle. Klin Wochenschr 4: 534–536

Warburg O, Minami S (1923) Versuche an Überlebendem Carcinom-gewebe. Klin Wochenschr 2: 776–777

Wicker CA, Hunt BG, Krishnan S, Aziz K, Parajuli S, Palackdharry S, Elaban WR, Wise-Draper TM, Mills GB, Waltz SE et al (2021) Glutaminase inhibition with telaglenastat (CB-839) improves treatment response in combination with ionizing radiation in head and neck squamous cell carcinoma models. Cancer Lett 502: 180–188

Yang W, Soares J, Greninger P, Edelman EJ, Lightfoot H, Forbes S, Bindal N, Beare D, Smith JA, Thompson IR et al (2012) Genomics of Drug Sensitivity in Cancer (GDSC): a resource for therapeutic biomarker discovery in cancer cells. Nucleic Acids Res 41: D955–D961

